# Ongoing introgression of a secondary sexual plumage trait in a stable avian hybrid zone

**DOI:** 10.1101/2023.03.30.535000

**Authors:** Kira M. Long, Angel G. Rivera-Colón, Kevin F. P. Bennett, Julian M. Catchen, Michael J. Braun, Jeffrey D. Brawn

## Abstract

Hybrid zones are dynamic systems where natural selection, sexual selection, and other evolutionary forces can act on reshuffled combinations of distinct genomes. The movement of hybrid zones, individual traits, or both are of particular interest for understanding the interplay between selective processes. In a hybrid zone involving two lek-breeding birds, secondary sexual plumage traits of *Manacus vitellinus*, including bright yellow collar and olive belly color, have introgressed asymmetrically ∼50 km across the genomic center of the zone into populations more genetically similar to *Manacus candei*. Males with yellow collars are preferred by females and are more aggressive than parental *M. candei*, suggesting that sexual selection was responsible for the introgression of male traits. We assessed the spatial and temporal dynamics of this hybrid zone using historical (1989 - 1994) and contemporary (2017 - 2020) transect samples to survey both morphological and genetic variation. Genome-wide SNP data and several male phenotypic traits show that the genomic center of the zone has remained spatially stable, whereas the olive belly color of male *M. vitellinus* has continued to introgress over this time period. Our data suggest that sexual selection can continue to shape phenotypes dynamically, independent of a stable genomic transition between species.

## Introduction

Hybridization is pervasive in nature and a key contributor to speciation and diversification processes (Abbott et al., 2013). Hybrid zones provide opportunities to explore how species originate from diverging populations that may still be experiencing some degree of gene flow. Several speciation and diversification processes can be observed in hybrid zones, including reinforcement of barriers to admixture (Coyne & Orr, 2004; Roberts & Mendelson, 2020), breakdown of reproductive barriers allowing fusion between hybridizing species (Kearns et al., 2018; Seehausen, Alphen, & Witte, 1997), creation of entirely new forms through hybrid speciation (Gross & Rieseberg, 2005), and adaptive introgression of traits from one species into another. Other hybrid zones appear to be at least quasi-stable, remaining in stasis for indefinite periods of time (Pinto, Titus-McQuillan, Daza, & Gamble, 2019; Wang, Rohwer, Delmore, & Irwin, 2019); yet closer study of such zones may reveal dynamic equilibria between opposing evolutionary forces (Barton, 1979; Barton & Hewitt, 1985).

The movement or stability of hybrid zones is influenced by many different evolutionary factors. Moving hybrid zones are observed in several taxa and can move slowly or many kilometers in a single year, especially if one parental species has a selective or competitive advantage over the other species (Dasmahapatra et al., 2002; Harr & Price, 2014; Metzler, Knief, Peñalba, & Wolf, 2021; Reudink, Mech, Mullen, & Curry, 2007; Taylor et al., 2014; Wang et al., 2014; Wielstra, Burke, Butlin, & Arntzen, 2017; Zohren et al., 2016). Conversely, prolonged spatial stability is often seen in “tension zones” (Barton & Hewitt, 1985) where hybrids have lower fitness than parentals and the zones are localized in specific regions coinciding with population density troughs or ecological transition zones (Barton & Hewitt, 1985; Rosser, Dasmahapatra, & Mallet, 2014; Ruegg, 2008; Smith, Hale, Kearney, Austin, & Melville, 2013). Thus, hybrid zones present dynamic systems governed by diverse factors, and their mobility or stasis may change over time along with changing evolutionary forces (Engebretsen, Barrow, Rittmeyer, Brown, & Lemmon, 2016; Wang et al., 2019). Moreover, hybrid zones often involve conflicting evolutionary forces, such as sexual selection and natural selection, where selection for the traits of one form can be opposed by generalized selection against hybrids. The outcome of these pressures over time and space is a topic of active investigation (Buggs, 2007; Wielstra, 2019; Yang et al., 2022). Characterizing phenotypic and/or genetic changes in hybrid zones over time can elucidate how species boundaries may go through periods of flux and allow us to capture the dynamics of evolution within these populations.

Genome-scale analyses now provide the power to parse evolutionary change in unprecedented detail, revealing cryptic gene flow, demographic and selective history, and the genetic underpinnings of key traits in speciation and adaptation (Abbott, Barton, & Good, 2016; Moran et al., 2021; Payseur & Rieseberg, 2016; Taylor & Larson, 2019). For example, evidence for the prevalence of ongoing gene flow during speciation has expanded dramatically with advances in genome sequencing (Chan et al., 2020; Trigo et al., 2013). Additionally, modern genomics allows for the detailed characterization of hybrid zone movement in real time and identification of loci responsible for introgressing traits (Billerman, Cicero, Bowie, & Carling, 2019; Ottenburghs et al., 2017; Semenov et al., 2021; Wielstra, 2019).

Hybrid systems with historical genetic sampling are particularly informative for understanding spatial dynamics in hybrid zones. One such system is a hybrid zone between golden-collared manakins (*Manacus vitellinus*) and white-collared manakins (*Manacus candei*) in western Panama. Both are lek breeding species, in which males compete for mating opportunities by performing elaborate displays (Day et al., 2021; Kirwan & Green, 2011), leading to strong sexual selection on male traits (Snow, 2004). This hybrid zone was first characterized nearly 30 years ago by studying clinal variation in morphological and genetic traits that differ between the parental forms along ∼100 km region of a 570 km transect (Brumfield, Jernigan, McDonald, & Braun, 2001; Parsons, Olson, & Braun, 1993). Male collar and belly color transition across this hybrid zone from a golden yellow throat and olive belly in the golden-collared manakin to a white throat and yellow belly in the white-collared manakin (Fig. 1) (Brumfield et al., 2001; Parsons et al., 1993). The phenotypic transition for collar color occurred at the Río Changuinola, the largest river in the region, where birds on one side of the river had yellow collars and birds on the other side had white collars (Parsons et al., 1993). This phenotypic transition was displaced by about 50 km from the cline centers for all genetic markers and several morphometric traits (Brumfield et al., 2001). A similarly displaced transition was observed for male belly color, although that cline was not as sharp as the collar color cline (Brumfield et al., 2001; Parsons et al., 1993). These displaced clines result in a geographic region where males resemble golden-collared manakins in two secondary sexual plumage traits, but are otherwise similar to white-collared manakins genetically and morphometrically (Brumfield et al., 2001; Parchman et al., 2013; Parsons et al., 1993). Assays of male aggression (McDonald, Clay, Brumfield, & Braun, 2001) and female choice (Stein & Uy, 2006b) indicated that yellow collars are advantageous over white collars, suggesting that sexual selection was responsible for collar color introgression. Belly color is correlated with collar color in these birds (Brumfield et al., 2001), and thus may be directly or indirectly under sexual selection as well.

**Figure 1.**
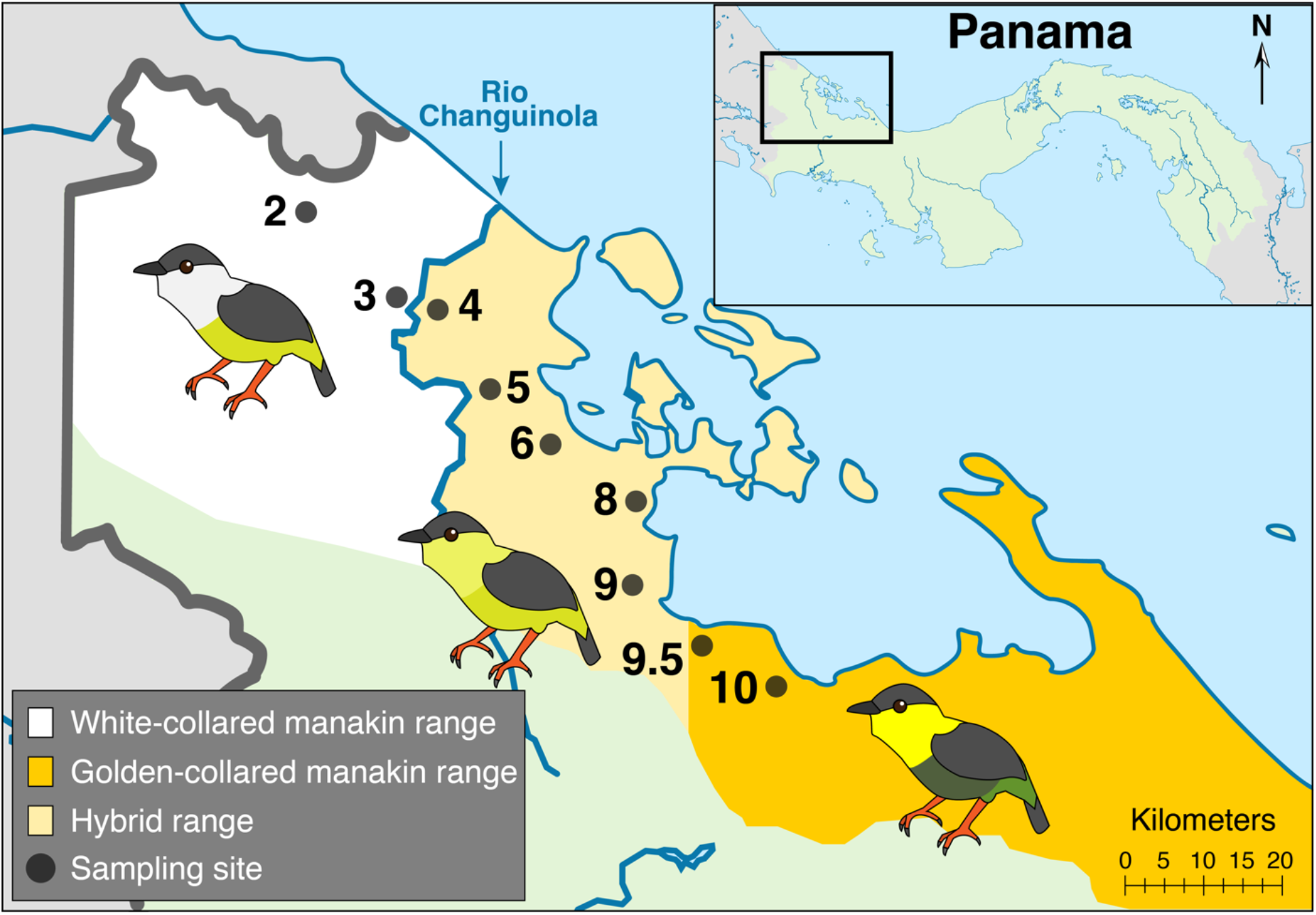
Map of the *Manacus* hybrid zone in Bocas del Toro, Panama. Dots represent sampling sites, numbered according to the system used in Brumfield et al. (2001). The shaded regions denote the approximate ranges of each parental species and hybrids in Panama based on the historical sampling. The cartoon birds illustrate the major phenotypic difference between the parental species and hybrid populations.

Our overarching aims are to understand the spatial and temporal dynamics of hybrid zones and how they may be influenced by a mating system that promotes strong sexual selection. We ask whether a strong preference for traits from one species over another can lead to the movement of an entire hybrid zone. Under this scenario, can particular traits become uncoupled from the rest of the genome? Do male secondary sexual plumage traits continue to introgress? We test these questions in a hybrid zone where strong selection for traits of one species have asymmetrically introgressed across the hybrid zone, but no information exists on the stability of the rest of the genome. Here, we report the results of a longitudinal reassessment of morphological and genetic variation across the *Manacus* hybrid zone using a newly collected set of contemporary (2017 - 2020) samples that closely replicates the historical (1989 - 1994) transect. We characterized the genetic structure and admixture of the hybrid zone in both the historical and contemporary transects using a genome-wide set of single nucleotide polymorphism (SNP) markers derived from Restriction site-Associated DNA sequencing (RADseq). We also estimated cline structure at both time points for previously identified male phenotypic traits of interest. We leveraged this multi-year sampling to explore the temporal dynamics of this hybrid zone, directly comparing the two datasets to provide empirical quantification for movement and stability in this hybrid system.

## Methods

### Study species and sampling sites

*Manacus vitellinus* and *M. candei* (Pipridae) are frugivorous birds that inhabit the understory of secondary forest or forest edge. *M. vitellinus* occurs from northern Colombia to Panama while *M. candei* ranges from southern Mexico to western Panama. For our historical dataset, we used many of the same specimens from the core transect collections studied by Parsons et al. (1993), Brumfield et al. (2001), Yuri et al. (2009), and Parchman et al. (2013). We then revisited the same sampling sites to collect contemporary samples. We replicated the core transect sampling design of Brumfield et al. (2001) as closely as possible, given land use changes in the intervening years between historical sampling in 1989-1994 and contemporary sampling in 2017-2020. We sampled *Manacus* populations at 9 distinct sites in Bocas del Toro, Panama, located an average of 11.56 km apart (Fig. 1). We used site numbers corresponding to the numbering scheme used in Brumfield et al. (2001). Sampling site 9.5 (Miramar), located between sites 9 and 10, was added to the contemporary dataset to increase the spatial resolution of geographic cline analysis (see methods below) and did not have previous historical transect data. We chose to focus our sampling within the active hybrid zone area, so sampling sites outside the province of Bocas del Toro were excluded, along with site 7 which deviated the farthest from the transect line. The 9 sampling sites represent two populations of *M. candei*, two populations of *M. vitellinus*, and 5 populations of hybrids in the known transition zone (Fig. 1, Table 1).

### Field and museum data collection

We used 36mm mesh “ATX” type mist nets on leks to capture individuals. Each bird was given a numbered aluminum band and a combination of plastic colored leg bands to facilitate individual identification. Birds were examined to determine age and sex using field observations of definitive adult male plumage and the presence of a brood patch for reproductive females.

**Table 1.** Sampling sites and number of sequenced individuals retained for genetic analyses after quality filtering.

| Sampling Site Number | Location Name | Distance from Site 10 (km) | Putative Hybrid Status | Historical N | Contemporary N |
| --- | --- | --- | --- | --- | --- |
| 2 | San San Drury | 92.50 | *M. candei* | 31 | 49 |
| 3 | El Silencio | 79.00 | *M. candei* | 19 | 26 |
| 4 | Finca Cuatro | 71.25 | Hybrid | 18 | 24 |
| 5 | Río Oeste | 48.50 | Hybrid | 17 | 20 |
| 6 | Quebrada Pastores | 42.50 | Hybrid | 4 | 24 |
| 8 | Río Uyama | 29.50 | Hybrid | 22 | 27 |
| 9 | Palma Real | 20.75 | Hybrid | 23 | 65 |
| 9.5 | Miramar | 8.44 | *M. vitellinus* | NA | 30 |
| 10 | Rambala | 0.00 | *M. vitellinus* | 18 | 53 |

Non-breeding females and juvenile males were sexed using a molecular sexing protocol (see below). We collected blood samples (<100 μL) in glass capillary tubes from all individuals by puncturing the brachial vein. Blood was stored in Longmire’s lysis buffer (Longmire, Maltbie, & Baker, 1997) and kept frozen to preserve long-term DNA quality. From wild-caught males with definitive adult plumage, we measured beard length (measured as the longest feather directly below the front of the eye), epaulet width (measured as the widest point of the white or yellow coverts from the bend of the folded wing to the posterior edge of the white or yellow epaulet feathers), and tail length (measured as the longest feather in the tail). These three morphological traits were identified as clinal in Brumfield et al. (2001). We remeasured these same three male phenotypic traits for the historical dataset on museum specimens at the Smithsonian National Museum of Natural History (NMNH). In order to minimize observer bias between the historical and contemporary datasets, all measurements reported here were taken by KML.

We characterized male plumage colors using Ocean Insight reflectance spectrophotometers (USB2000+ in 2017 and FLAME-S-UV-VIS-ES in 2018-2022). Both spectrophotometers used a PX-2 pulsed xenon lamp (Ocean Insight). We used a fiber optic probe fitted with a plastic probe holder and took the feather color reads at a 90° angle. The probe holder minimized the ambient light in reflectance measurements and maintained the same distance between the probe and the feathers for each read. The percent reflectance across all wavelengths was measured relative to a pure white reflecting standard (WS-1-SL Diffuse Reflectance Std, Spectralon) as 100% reflectance and a 0% dark standard. The white standard was fitted with a plastic holder that kept the probe from directly touching the white substrate of the standard, keeping the white standard clean and free of discolorations. Due to the added distance between the probe holder and the white standard substrate, we calibrated our measurements to account for the drop in reflectance from this gap, which would be present evenly across the entire spectrum on each spectrophotometer. We adjusted the spectral curve values using a 0.7 and 0.5 multiplier for the USB2000+ and FLAME-S-UV-VIS-ES spectrophotometers, respectively, based on the percent drop in white reflectance as measured on both spectrophotometers following the calibration methods of Cook et al. (2013), Kelly et al. (2012), and Troy Murphy (personal communication). To assess bias in the data due to using two different spectrophotometers, we plotted the reads taken by each spectrophotometer and did not observe shifts in reads based on which spectrophotometer was used (Fig. S1, Table S1).

The reflectance data for the contemporary birds were taken on live birds in the field, while the reflectance data for the historical specimens were measured from museum skins at the Smithsonian NMNH using the FLAME-S-UV-VIS-ES spectrophotometer. Previous studies have found that the plumage of museum specimens can change over time but that, overall, the reflectance spectra from well-preserved museum specimens are similar to wild birds (Doucet & Hill 2009). Additionally, museum specimen degradation will depend on how well preserved the specimens are, the age of the specimen, and the mechanism underlying the color of interest. The *Manacus* specimens at the Smithsonian NMNH used in this study are between 31 and 26 years old and are clean and in good condition. We took reflectance spectra of definitive male collar and belly plumage color near the center of the throat or belly on each bird. Five spectra were taken for each plumage patch, repositioning the probe between each read and averaging across spectra for each bird. Following Brumfield et al (2001), we used the average adjusted reflectance at 490 nm and 665 nm to represent collar color and belly color respectively, because these wavelengths had the greatest differences in mean reflectance between the two parental samples. Outlier readings beyond two standard deviations (SDs) away from the mean for any individual were removed from the dataset. We used the average percent reflectance at each wavelength for geographic cline analysis.

All protocols involving live birds were reviewed and approved by the Illinois Animal Care and Use Committee and the Smithsonian Tropical Research Institute Animal Care and Use Committee (Illinois IACUC numbers: 15234 & 18239; STRI ACUC numbers: 2016-0301-2019 & 2019-0115-2022).

### RADseq library preparation and genotyping

DNAs from the contemporary 2017-2020 samples were extracted from whole blood using the animal tissue protocol on a Gene Prep automated extractor (Autogen), which involves Proteinase K digestion, phenol extraction, and alcohol precipitation. DNA concentrations were determined using a Quant-iT™ dsDNA Broad-Range Assay Kit (Thermo Fisher Scientific) on a SpectraMax® iD3 plate reader running SoftMax® Pro 7 software. DNA stocks for the historical 1989-1994 samples were available from previous studies (Brumfield et al., 2001; Parsons et al., 1993). All samples are deposited at the National Museum of Natural History, Smithsonian Institution, in Washington, D.C. DNA from individuals of unknown sex were assayed in an avian polymerase chain reaction (PCR) molecular sexing protocol (Fridolfsson & Ellegren, 1999). Briefly, we made a master mix using the primers 2550F = 5’-GTT ACT GAT TCG TCT ACG AGA-3’ and 2718R = 5’-ATT GAA ATG ATC CAG TGC TTG-3’ and TaKaRa ExTaq DNA Polymerase (Takara Bio USA, Inc). We ran the samples for 31 cycles and ran the PCR product on a 2% agarose gel.

Six single-digest *SbfI* RADseq libraries were prepared using the protocols described in Baird et al. (2008) and Etter et al. (2011). After diluting all samples in the library to the same concentration, 600-800 ng of genomic DNA per-sample (depending on the library, see Table S2, Table S3) was digested using the *SbfI*-HF enzyme (New England Biolabs). Custom P1 adapters containing unique 7-bp barcodes (Hohenlohe, Bassham, Currey, & Cresko, 2012) were ligated to each individual sample with T4 ligase (New England Biolabs). The DNA from all uniquely barcoded individuals in a library was pooled at equimolar concentration, and 1500 ng of total DNA was then sheared using Covaris (Covaris, Inc.), or Qsonica (Qsonica LLC) sonicators and size selected for 300-600 bp inserts using AMPure XP beads (Beckman Coulter). The sheared DNA was end-repaired and ligated to custom P2 adapters. The P2-ligated DNA was then amplified for 12 rounds of PCR using custom RAD primers (Hohenlohe et al., 2012) and cleaned with 0.8X AMPure XP beads (Beckman Coulter). The final libraries were sequenced either on an Illumina HiSeq 4000 or an Illumina NovaSeq 6000 to generate 2×150 bp reads (Table S3).

In total, 593 samples were processed, which included 542 unique individuals and 51 samples sequenced in duplicate to ameliorate low coverage. A breakdown of the detailed sequencing information per-library is provided in Table S3. Using the *Stacks* software version 2.60 (Rochette, Rivera-Colón, & Catchen, 2019), raw reads were filtered with `process_radtags` to remove reads with low quality (`-q`) and uncalled bases (`-c`), rescue ambiguous barcodes (`-r`), and demultiplex samples based on their unique barcode (`-b`). For each library, we retained between 668 million to 1.6 billion reads (Table S3).

The processed reads for all 593 sequenced samples were mapped to the *M. vitellinus* reference assembly (NCBI RefSeq Accession: GCF_001715985.3) using *BWA* `mem` version 0.1.17 (H. Li & Durbin, 2009; Heng Li, 2013) with default parameters. Alignments were processed and sorted using *SAMtools* version 1.7 `view` and `sort` (H. Li et al., 2009). RAD loci assembly and genotyping from the processed alignments was done with the *Stacks* `gstacks` module version 2.60 (Rochette et al., 2019), where we removed PCR duplicates (`--rm-pcr-duplicates`). In total, we assembled 426,521 raw RAD loci with an average non-redundant coverage of 14.0× (SD 6.9×). To provide a baseline level filtering of missing data, we ran the *Stacks* `populations` module on all samples with an average per-locus coverage at 6× or higher, requiring a minimum minor allele count of 3 (`--mac 3`), requiring a locus to be present in at least 80% of samples per population (i.e., sampling sites) (`-r 0.80`), and be in all 9 populations (`-p 9`). Additionally, we used the `--hwe` flag to calculate differences between the observed and expected genotype frequencies under Hardy-Weinberg equilibrium (HWE) for each population. This filtered run kept 77,551 loci and 584,680 variant sites across the 470 individuals that were retained in the dataset (Table 1). This `populations` run is hereafter referred to as the “master run” and its output was used to create all whitelists of loci/SNPs for downstream analyses.

### Population structure analysis

We used *ADMIXTURE* (Alexander, Novembre, & Lange, 2009) to assess population genetic structure in both the historical and contemporary datasets. Using a custom Python script, we created a whitelist of 10,000 SNPs sampled from our master `populations` run. First, we filtered the data from the master run for SNPs with a minimum allele frequency (MAF) of 1% and in HWE in all populations, and selected a single, randomly chosen SNP at each locus in order to ensure the independence of markers in the dataset. From this filtered subset, the script then randomly selects 10,000 sites to generate a final whitelist. For each dataset, we reran the *Stacks* `populations` module, supplying the whitelist and exporting genotypes in the *PLINK* (`--plink`) and *GENEPOP* (`--genepop`) format. The resulting `.ped` files were converted to binary `.bed` files using *PLINK* version 1.90b6.25 (Chang et al., 2015) and *ADMIXTURE* version 1.3.0 was run separately on the historical and contemporary datasets for population cluster (K) values 1-9. To find an optimal value for the number of population clusters (K), we followed the *ADMIXTURE* documentation by plotting the cross validation (CV) error for each K value. To corroborate the results of *ADMIXTURE* using an ordination method instead of a clustering method, we ran a principal component analysis (PCA) using the same whitelisted SNP dataset described above in the *ADMIXTURE* analysis. We ran the PCA in *adegenet* version 2.1.7, an *R* package for multivariate analysis of genetic markers (Jombart, 2008; Jombart & Ahmed, 2011). The *ADMIXTURE* and PCA results were plotted in *R* version 4.2.1 (R Core Team, 2022).

### Hybrid index

In order to compare the shifts the location of the genetic center of the hybrid zone over time, we calculated the hybrid index for each of 470 individuals across historical and contemporary datasets using the *R* package *gghybrid* version 1.0.0 (Bailey, 2020), which uses a Bayesian Markov chain Monte Carlo (MCMC) method for estimating the hybrid index. First, using the same whitelist of 10,000 SNPs as the *ADMIXTURE* analysis, we re-ran `populations` to export the genotypes in the *STRUCTURE* format (`--structure`). Using the *gghybrid* data input function (`data.prep()`), we assigned the parental source population 0 and 1 as *M. candei* (sampling site 2) and *M. vitellinus* (sampling site 10), respectively. Using the `max.S.MAF` option, we removed loci for which the smaller of the two parental minor allele frequencies is greater than 10%, ensuring that all SNPs used in the hybrid index are informative by requiring the allele frequency of at least one parental population to be close to 0. This filter also allows for the retention of SNPs that are not fixed for alternate alleles in parental populations, providing for denser sampling of the entire genome in estimating the hybrid index. We calculated the Bayesian hybrid index using the `esth()` *gghybrid* function, running for a total of 50,000 iterations with 10,000 burn-ins. This analysis was performed separately for the historical and contemporary datasets. We recorded the mean hybrid index value for each individual and used the mean for each sampling site in downstream analyses.

### Assignment of hybrid class

We classified our 470 *Manacus* individuals among different hybrid types (i.e., if an individual is an F1, F2, backcross, etc.) by establishing the relationship between the parental proportion (in the form of hybrid index) and their interspecific heterozygosity. The interspecific heterozygosity statistic calculates the proportion of an individual’s genome that contains alleles inherited from both parental populations. To establish this relationship, we first obtained a subset of 2,000 diagnostic alleles between the two parental populations (see supplemental methods). Diagnostic alleles were used as the estimation of interspecific heterozygosity can be sensitive to alleles of intermediate frequencies present in the parental populations (Fitzpatrick, 2012).

Using the genotypes of these 2,000 diagnostic markers, we first re-estimated the hybrid index of the 470 individuals using the methods described above (using the `esth()` function in *gghybrid*). Then, we calculated interspecific heterozygosity using the R package *Introgress* version 1.2.3 (Gompert & Buerkle, 2010). In *Introgress*, we first processed the genotype data for both parental and admixed individuals using the `prepare.data()` function, defining the input genotypes as co-dominant markers (coded as “`C`” in the `loci.data` file) and not fixed in the parental populations (`fixed=FALSE`). The interspecies heterozygosity of each individual was then estimated using the `calc.intersp.het()` function. We used “triangle plots” to show the interspecific heterozygosity as a function of hybrid index (Fig. S4), classifying individuals among different hybrid types based on their placement between the two axes.

The hybrid classification analysis was additionally repeated on a simulated dataset. Multi-generation hybrid genotypes were simulated from the pool of parental genotypes obtained from the 2,000 diagnostic SNPs. Our simulations were performed using a custom Python script (see supplemental methods), using a similar approach to the methods described by (Lavretsky, Janzen, & McCracken, 2019; Lavretsky et al., 2016; Wringe, Stanley, Jeffery, Anderson, & Bradbury, 2017). Briefly, we used the empirical parental genotypes to calculate the observed allele frequency across our 2,000 diagnostic SNPs. Then, we defined a series of defined crosses, each describing the generations of hybridization (or backcross), the parents of each cross, as well as the genomic proportions (Fitzpatrick, 2012; Turelli & Orr, 2000) expected. Lastly, the parental allele frequencies and genomic proportions were then used to determine the probability of genotypes in the individuals of a given hybrid cross. The genotypes of simulated individuals of known hybrid assignment were then used to calculate hybrid index and interspecific heterozygosity (Fig. S4), which were then compared against the values observed in our *Manacus* empirical data.

### Geographic clines

For the geographic clines, we re-filtered the master dataset to obtain all sites that were in HWE in each of the 9 populations and only a single SNP per locus, retaining a total of 72,506 loci. Sites in HWE were used for the geographic cline analysis because sites that drastically depart from equilibrium could bias cline fittings (Derryberry, Derryberry, Maley, & Brumfield, 2014; Macholán et al., 2008, 2007; Phillips, Baird, & Moritz, 2004). All geographic cline analyses were repeated with and without the newly added sampling site 9.5. With the new 72.5K SNP whitelist, we re-ran the *Stacks* `populations` module to export genotypes in the *HZAR* format (`--hzar`). *HZAR* is an *R* package designed to fit geographic clines to one-dimensional transect data in hybrid zones using the Metropolis-Hastings Markov chain Monte Carlo algorithm (Derryberry et al., 2014). We wrote a custom *R* script to parallelize *HZAR* version 0.2-5, allowing us to efficiently run the analysis independently for each selected SNP. Of the 72.5K whitelisted markers, we removed sites that were invariant in either the contemporary or historical datasets, ensuring that only sites that were variant in both datasets were run in *HZAR*. In total, we retained 68,684 and 63,444 SNPs in the contemporary and historical datasets, respectively. This relatively large number of markers was retained for each dataset in order to maximize sampling of clinal variation across the genome.

For each SNP in the historical and contemporary datasets, we ran *HZAR* using four cline models (as originally described by (Szymura & Barton, 1986, 1991)) to maintain consistency with the analyses performed by Brumfield et al. (2001). The four models were: 1) a model with the observed minimum (*p*_min_) and maximum (*p*_max_) allele frequencies and without fitting exponential decay curves for the cline tails, 2) a model with estimated *p*_min_ and *p*_max_ without fitted exponential decay curves, 3) a model with estimated *p*_min_ and *p*_max_ and fitting decay curves to both cline tails, and 4) a null model without fit for either allele frequency or cline tails. For each model, we ran 3 independent chains, composed of three dependent model fit requests, each of length 100,000 and with 10,000 burn-in generations. We randomly selected a subset of the genetic clines to inspect the MCMC trace plots of the chains to verify that the models were converging to a local optimum. The best fit model was selected using Akaike Information Criterion with correction for small sample size (AICc). We then extracted the estimated cline width and cline center values with 95% Bayesian Credibility Intervals (CI) for the best model of each cline, along with the maximum likelihood cline generated by the best model. We removed sites that were assigned null models (i.e., indicating that the SNP was not clinal and thus contained null values for the cline center and width). In the historical and contemporary datasets, there were 11,925 and 8,254 sites, respectively, that were non-clinal and thus were assigned null models. We used a custom Python script to combine the cline parameter outputs into a single file and to select clines at loci that were present in both the historical and contemporary datasets. The distribution of all 48,316 cline centers is shown in Fig. S2. Next, we removed SNPs displaying cline centers located beyond the geographical boundaries of the studied area (i.e., cline centers less than 0 Km or greater than 92.5 Km). In total, we retained cline results for 46,734 SNPs that passed all the above filters and were observed in both historical and contemporary datasets (comprising our base filtering scheme). Additionally, we applied subsequent filters based on the difference in allele frequencies between the parental sampling sites and the range of the estimated CI, to further validate our detection of movement between temporal datasets (see supplemental methods). We considered a cline as showing evidence of movement if the cline center CI did not overlap between historical and contemporary datasets.

We also ran *HZAR* on five male phenotypic traits known to differ between *M. vitellinus* and *M. candei*: beard length, epaulet width, tail length, collar color, and belly color. We ran three morphological cline models: 1) a model with the observed morphological data (“fixed”) and without fitting exponential decay curves for the cline tails, 2) a model with estimated “free” without fitted exponential decay curves, 3) a model with estimated “free” and fitting decay curves to both cline tails. After running each model three times with a chain length of 100,000, and burn-in of 10,000, the best fit model was selected using AICc. The cline analyses for the contemporary dataset were performed both with and without data for site 9.5 to directly match the historical dataset. For any phenotypic traits that displayed substantial changes in cline center or width estimates, we further validated these results by running longer chain lengths of 1,000,000 with 100,000 burn-in, and visually inspecting the MCMC trace plots. In addition to the five male phenotype clines, we ran a geographic cline on the genomic hybrid index calculated by *gghybrid*, to assess genome-wide movement of the hybrid zone. We calculated the mean hybrid index values in each sampling site for both the contemporary and historical datasets and analyzed them using *HZAR*. As with the SNP data, we obtained the best cline model and exported the associated center, width, and CI values. For both the phenotypic clines and hybrid index clines, we assessed whether the clines had moved based on whether the historical and contemporary CIs for the cline centers overlapped.

## Results

### Genome-wide patterns of genetic population structure are stable through time

We sampled 10,000 SNPs from 470 sequenced individuals spanning the historical and contemporary datasets for admixture analyses. Overall, we found genetic population structure across the transect to be consistent over time (Fig. 2A). When we assessed admixture with two genetic clusters (K=2), individuals separated based on apparent parental species ancestry. The genomic transition from *M. vitellinus* to *M. candei* was centered near site 9, with substantially admixed individuals present only at sites 8, 9 and 9.5. Although all birds in sites 4-8 closely resemble *vitellinus* phenotypically, their genomes are *candei*-like, aside from a few intermediate individuals at site 8.

**Figure 2.**
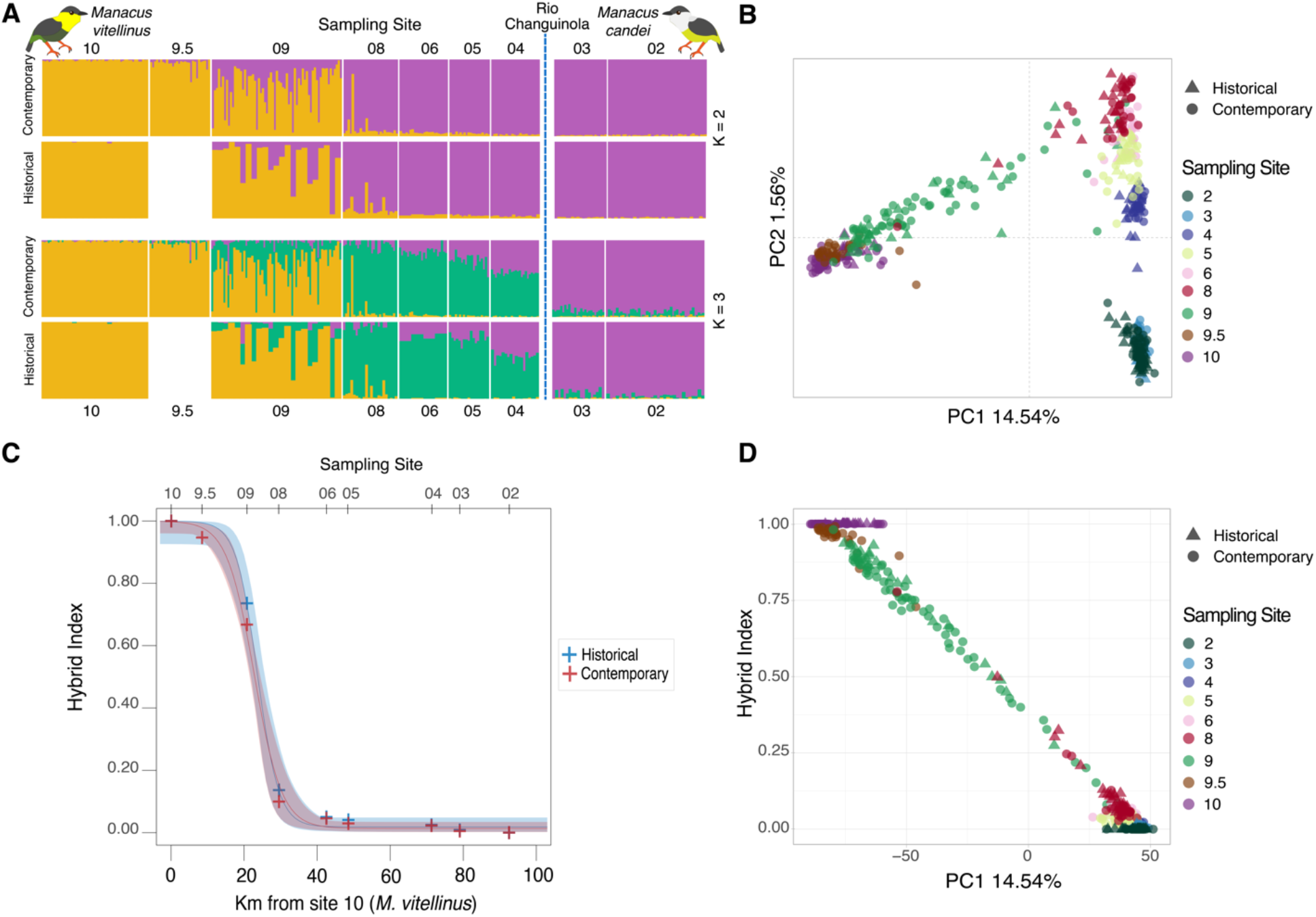
Patterns of population structure across the *Manacus* hybrid zone. **A)** Admixture analysis of contemporary and historical manakin datasets based on 10,000 SNPs. Sampling site numbers are listed at the top and bottom of the figure with the *M. vitellinus* site 10 on the left and *M. candei* site 2 on the right. Site 9.5 was sampled for the contemporary but not the historical dataset. Each vertical bar represents one bird; width of contemporary and historical population blocks was equalized for ease of comparison. The dashed blue vertical line between sites 3 and 4 represents the Río Changuinola, marking the phenotypic transition of yellow to white collar plumage. **B)** Principal component analysis based on 10,000 SNPs from contemporary and historical manakin datasets. **C)** Geographic clines of the hybrid index and associated 95% credibility intervals, with the historical dataset in blue and contemporary in red. A hybrid index of 0 indicates complete *M. candei* ancestry (site 2) while a value of 1 indicates complete *M. vitellinus* ancestry (site 10). The top x-axis shows the location of the 9 sampling sites. The crosses denote the mean hybrid index at each site. **D)** Correlation between the first principal component and genomic hybrid index.

K=3 was the best fit model for both historical and contemporary datasets according to cross validation analyses (Table S4). With K=3, a third genetic cluster separated hybrids with candei-like genomes east of the Río Changuinola (sites 4-9) from *M. candei* to the west (sites 2 and 3), highlighting the river as an important barrier to gene flow (Fig. 2A). The largest variance in admixture proportions is still seen in sampling site 9 in both time points. Patterns of admixture are consistent at K=2 and K=3 clustering for the historical and contemporary transect samples, showing that overall admixture and genetic clustering in the hybrid zone is stable over the time period examined.

The PCA corroborated the admixture analysis (Fig. 2B). Individuals in the PCA group by sampling site and not by time, also implying temporal stability in genome-wide structure. The first principal component axis (PC1) separates individuals based on apparent parental ancestry, explaining 14.54% of SNP variance. We found a tight correlation between PC1 scores and genomic hybrid index, suggesting that the primary axis of SNP variation strongly corresponds with parental ancestry (Fig. 2D). Sampling site 9, again, showed the largest variation of all the sampling sites, spanning between the *M. vitellinus* parental cluster and the other hybrid sites. Additionally, parental *M. candei* sites 2 and 3, both to the west of the Río Changuinola, clustered tightly together and are separated from the rest of the hybrid sites along PC2; however, this axis only explains a small proportion of the observed variation (1.56%).

### Genomic hybrid index reflects stable patterns of parental ancestry

The distributions of hybrid indices were stable over time and showed no substantial movement of the principal genomic transition in the hybrid zone (Fig. 2C). The geographic clines for the genomic hybrid index had very similar cline centers and cline widths with largely overlapping credibility intervals from the historical and contemporary transect datasets (Table S5). The cline center credibility intervals from both datasets overlapped the location of hybrid sampling site 9, about 20.75 km from *M. vitellinus* parental site 10. Thus, the genomic center of the hybrid zone has remained stable over time near sampling site 9.

Birds sampled at sites 4-8 for both contemporary and historical transects possessed a minor amount of *M. vitellinus* ancestry in a majority *M. candei* genomic background, with the exception of a single individual in contemporary site 8 with a hybrid index of 0.78 and a single individual in historical site 8 with a hybrid index of 0.50 (Fig S3, Table S6). The contemporary samples from newly-assayed site 9.5 showed a majority *M. vitellinus* ancestry, with a mean hybrid index of 0.9470 (SD 0.0612). While extensive admixture was observed at site 9 in both the contemporary and historical datasets, we did not find evidence of early generation hybrids in either the contemporary or historical samples (Fig. S4). Hybrids from site 9 had the highest levels of variation in hybrid index and interspecific heterozygosity in both the contemporary and historical datasets (Fig. S3, Fig. S4, Table S6). This result is consistent with the population level structure analyses above, showing that site 9 is consistently the most variable and admixed locality in the hybrid zone. Thus, the proportion of genome-wide parental ancestry in hybrid individuals appears stable over time at the population level.

### Genome-wide geographic clines have remained stable

Of the 72,506 SNPs in the full RADseq dataset, we retained 46,734 SNPs that displayed clinal variation across the transect and passed all baseline filters for the cline center analysis. The overall distribution of contemporary clines with and without the newly added sampling site 9.5 are shown in Fig. S2, but the inclusion of the additional site did not largely change the cline center distribution. The clines from the historical samples had a median cline center at 23.85 km from site 10, the *M. vitellinus* parental site (mean = 32.98 km, SD = 21.80 km). The contemporary dataset had a median cline center at 22.75 km from site 10 (mean = 30.81 km, SD = 19.41 km). The average cline center for both datasets appear to be largely robust to the effects of filtering, with the median cline centers estimated between 22.44 and 23.85 km for both datasets across all filtering schemes (Table S7).

Both the historical and contemporary median cline centers are between sampling sites 8 and 9, located at 29.50 km and 20.75 km from site 10, respectively. Similarly, 51.25% and 50.42% of all contemporary and historical clines, respectively, have estimated cline centers between these two sampling sites. While the specific proportion of cline centers located between these two sampling sites varies depending on the filters applied (ranging from 50.42 to 97.95%; Table S7), all filtering schemas find the majority of cline centers for the historical and contemporary datasets located in the 8.75 km between sampling sites 8 and 9. This distribution of cline centers surrounding sampling site 9 provides support for the assignment of this site as the genomic center of the *Manacus* hybrid zone, in accordance with previous findings.

While, on average, cline centers are located proximally to the previously characterized hybrid center (sampling site 9); we do find evidence of displaced clines where the shift in allele frequency occurs near the Río Changuinola (75.13 km away from sampling site 10). Under the base filtering scheme, we detected 3,166 (6.77%) clines with maximum likelihood cline center estimates at or past the river in the contemporary dataset, and 4,421 (9.46%) in the historical dataset. While the specific proportion of clines with centers at or past the river is dependent on the effects of filtering (ranging from 0 to 9.46%; Table S7), we do detect the displacement of a small, but measurable, proportion of clines away from the genetic center of the hybrid zone. For both the historical and contemporary datasets, these clines with centers along the Río Changuinola are likely examples of loci introgressing along the hybrid zone, shifting from the genetic hybrid center towards the current phenotypic transition.

We detected only 795 (1.70%) out of our total 46 thousand SNPs from our base filtering scheme exhibited evidence of movement over time, showing non-overlapping cline center CIs between the two temporal datasets. The proportion of clines centers with non-overlapping CIs ranged between 0.77% and 3.66% across all filtering schemas (Table S7). From the base filtering scheme, the 795 putatively moving clines are distributed across 91 different scaffolds in the *M. vitellinus* reference assembly (Table S8). At our most stringent filtering scheme, filtering by 80% allele frequency differences between the two parental populations to observe a subset of more putatively diagnostic alleles (see supplemental methods), we detect only six clines with evidence of movement located in six different scaffolds (Table S7 and Table S8). Since the *M. vitellinus* reference assembly is not chromosome-level, we used conserved synteny to identify orthologous chromosomes with the lance-tailed manakin (*Chiroxiphia lancaeolata*) reference assembly (see supplemental methods) to verify that the SNPs with evidence of movement are on several separate chromosomes and not clustered in a single region of the genome (Table S8). These results indicate that genomic loci have remained geographically stable over thirty years, but that limited cline movement has occurred.

### Plumage introgression has advanced in thirty years

In addition to the distribution of allele frequencies across space, we also fit sigmoid clines to the observed variation in five male plumage traits (beard length, epaulet width, tail length, collar color, and belly color) in both historical and contemporary transect samples (Table S9, Table S10). Clines for four plumage traits are similar in the historical and contemporary transects (Fig. 3A-D, Table S5), suggesting these traits are largely geographically stable through time. Beard length and tail length cline centers lie near the genomic center of the hybrid zone (site 9) at ∼23 km from sampling site 10, while the collar color cline centers lie more than 50 km further along the transect near the Río Changuinola (Fig. 3F). The historical and contemporary CIs for beard length cline centers overlap, but the CIs for cline widths do not, indicating that the cline has widened by ∼11 km (Table S5). This widening could indicate some degree of bidirectional introgression or relaxed selection on beard length, but we do not currently have further evidence to explain this change. Epaulet width is a shallow cline that spans the whole hybrid zone and does not exhibit a strong sigmoidal shape (Fig. 3B). Cline estimates for epaulet width were inconsistent across the analysis, with the best cline model switching between model I and III depending on the chain length and the addition of the contemporary sampling site 9.5 (Table S5). Given these results, we cannot accurately resolve any temporal changes (or lack thereof) in this trait.

**Figure 3.**
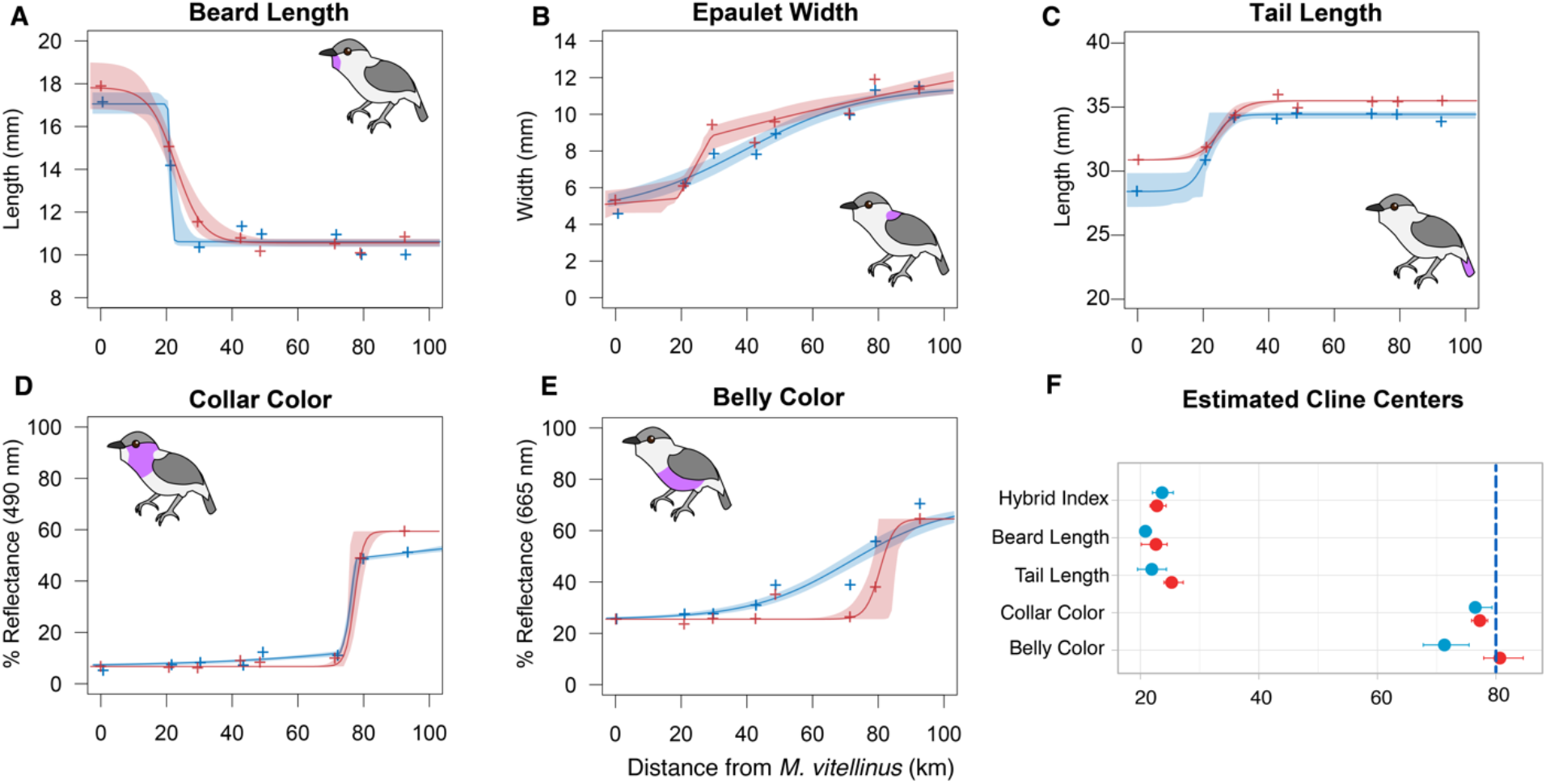
Contemporary and historical clines estimated from manakin phenotypic data across 8 sampling sites. For all panels, blue denotes the historical dataset, red the contemporary dataset, and all x-axes are in kilometers from the parental *M. vitellinus* (site 10). Crosses in **A-E** represent the average value of the phenotypic trait of interest in each sampling site. Shaded regions represent 95% credibility intervals for each cline. Magenta patches on the manakin cartoons highlight the plumage patches measured for each trait. **F)** Location along the transect of the maximum likelihood estimated cline center (dot) with 95% credibility intervals (horizontal bars) for the traits of interest. Hybrid index values pertain to the genomic hybrid index cline shown in Figure 2C. The vertical blue dashed line represents the location of the Río Changuinola.

In contrast, the olive belly coloration exhibits evidence of movement across this hybrid zone over the last 30 years (Fig. 3E, Table S5). The historical dataset has an estimated cline center for belly color at 71.31 km (95% CI 67.74 – 75.47 km) while the contemporary dataset has an estimated cline center at 80.47 km (95% CI 77.89 – 84.59 km). Based on non-overlapping CIs, we conclude that this cline has moved about 9 km in the last 30 years. The cline for belly color also narrowed significantly, with an estimated historical width of 60.58 km (95% CI 50.98 – 72.46 km) compared to only 8.73 km (95% CI 1.55 – 17.02 km) for the contemporary dataset.

The difference in the contemporary and historical belly color clines was consistent whether or not sampling site 9.5 was included in the analyses (Table S5). Thus, the belly color cline has shifted to the Río Changuinola (Fig. 3F) and is now similar in position and width to the collar color cline.

## Discussion

The hybrid zone between the golden-collared manakin and the white-collared manakin is one of the few documented cases showing asymmetrical introgression of a male secondary sexual trait. In agreement with previous research on this system (Brumfield et al., 2001; Parchman et al., 2013; Parsons et al., 1993), we see that yellow collars, which have been shown to be under positive sexual selection in this hybrid zone (McDonald et al., 2001; Stein & Uy, 2006b, 2006a), have been displaced up to the large regional river and define the phenotypic transition between the parental forms. We also found evidence of the displacement of several genetic clines toward this phenotypic transition, implying that there is historical introgression across several independent loci throughout the genome. Additionally, we found evidence that belly plumage coloration, a male secondary sexual trait, has moved and narrowed since the initial sampling, with hybrid males in the introgression zone getting darker green bellies over time. Although the possibility of direct sexual selection on belly color has not been formally tested, dark bellies are following the same introgression pattern as male yellow collar color. Together, these results suggest that dark bellies are being selected for in addition to yellow collars.

Despite this movement and displacement of putatively sexually selected traits and loci, we found clear evidence that the genomic transition of this hybrid zone and several phenotypic clines have remained spatially stable over the span of ∼25-30 years. The contrast between genomic stability and ongoing introgression of a putatively sexually selected trait implies that, while sexual selection is driving movement in several traits and genetic loci towards the river, there is a counteractive selective force maintaining the large-scale stability of the hybrid zone. While the specific mechanisms maintaining the stability of the hybrid zone are currently unknown, one counteracting force could be generalized selection against hybrid individuals through, for example, reduced reproductive success (Svedin, Wiley, Veen, Gustafsson, & Qvarnström, 2008) and/or genetic incompatibilities (Schumer et al., 2018). Independent of the specific mechanism of selection, our work highlights how the interplay of sexual selection and natural selection each shape the genomes of hybrids and the dynamics of the hybrid zone as a whole.

### Asymmetrical introgression in avian tension zones

Previous studies characterizing the *Manacus* hybrid zone phenotypically and genetically found evidence of asymmetrical introgression of male *M. vitellinus* plumage traits, yellow collar color and olive belly color, into populations dominated genetically by *M. candei* (Brumfield et al., 2001; Parchman et al., 2013; Parsons et al., 1993). Despite this introgression, we found that the *Manacus* hybrid zone remains narrow, consistent with previous studies which found 6 out of 7 genetic markers had cline widths of less than 11 km (Brumfield et al., 2001). Additionally, our genomic hybrid index and admixture analysis corroborate the previous studies (Brumfield et al., 2001; Parchman et al., 2013; Parsons et al., 1993), finding that sampling sites 4 - 8 are majority *M. candei*-like and that site 9 is the genomic hybrid center of the hybrid zone. The results present a clear picture of very limited genetic introgression across a geographically narrow, temporally stable hybrid zone.

Studies of other avian hybrid zones also report asymmetrical introgression within narrow hybrid zones, e.g., northern flickers (Aguillon & Rohwer, 2022); red-backed fairy-wrens (Baldassarre, White, Karubian, & Webster, 2014); long-tailed finch (Griffith & Hooper, 2017); wagtails (Semenov et al., 2021); jacanas (Lipshutz et al., 2019); tanagers (Morales-Rozo, Tenorio, Carling, & Cadena, 2017). Many of these hybrid zone studies posit that movement may be controlled by changes in the environment (Aguillon & Rohwer, 2022; Carling & Brumfield, 2008; Carling & Zuckerberg, 2011; Walsh, Billerman, Rohwer, Butcher, & Lovette, 2020), or climate change (Taylor et al., 2014); however, sexual selection appears to play a more important role in *Manacus*. Our results better align with other hybrid zones which show a trait under sexual selection displaced from the genomic hybrid center due to either intersexual (mate choice) or intrasexual (competition for access to mates) selection (Baldassarre et al., 2014; Lipshutz et al., 2019; Yang et al., 2018). The *Manacus* hybrid zone is a prime example of sexual selection driving key diagnostic traits across species boundaries; however, not all displaced clines represent ongoing introgression (Semenov et al., 2021). As such, reassessing hybrid zones with asymmetrical introgression over multiple time points helps to clarify which traits have ongoing introgression.

### Barriers to gene flow despite positive selection

Male collar color displayed nearly identical clines in the contemporary and historical datasets; specifically, a sharp transition in collar color from yellow to white occurring at the Río Changuinola (Fig. 1). Collar color likely introgressed through positive sexual selection mediated by female choice (Stein & Uy, 2006b), male aggression (McDonald et al., 2001), or both. If positive sexual selection is still operating, as it appears to be for belly color, collar color should continue to spread. However, yellow collar plumage appears to be “stuck” at the river, displaced by more than 50 km from the genomic transition, consistent with previous studies (Brumfield et al., 2001; Parchman et al., 2013; Parsons et al., 1993). Thus, the Río Changuinola appears to present a limit to collar color introgression, either by acting as a barrier to gene flow or marking a transition in another relevant evolutionary factor (Bennett, Lim, & Braun, 2021). Belly color introgression may prove to be similarly limited at the river.

Rivers have been shown as effective barriers to dispersal in tropical birds (Haffer, 1997; Moncrieff et al., 2024; Naka, Bechtoldt, Henriques, & Brumfield, 2012; Naka, Costa, Lima, & Claramunt, 2022) and other taxa, e.g., frogs (Fouquet et al. 2012), primates (Fordham, Shanee, & Peck, 2020), and butterflies (Rosser, Shirai, Dasmahapatra, Mallet, & Freitas, 2021). However, the extent that a river, or any geographic barrier, serves to reduce gene flow likely depends on the dispersal mode or capability of the species of interest (Claramunt, Derryberry, Remsen, & Brumfield, 2012; Claramunt, Hong, & Bravo, 2022; Naka et al., 2022; Nazareno, Knowles, Dick, & Lohmann, 2021). The golden-collared manakin is capable of flying substantial distances over open water (Moore, Robinson, Lovette, & Robinson, 2008), and we observed radio-tagged females dispersing beyond a kilometer from the lek where they were tagged. In one case, a female *M. vitellinus* individual was tracked flying about 100m over open cattle pasture to a neighboring stand of trees to forage before flying back to the lek in an isolated forest patch (KML personal observation). Additionally, *Manacus* manakins are secondary forest specialists and most leks in our study area are along waterways or flanked by cacao plantations or cattle pastures.

Thus, while the Río Changuinola seems to be acting as a barrier to gene flow given the location of the displaced clines, *Manacus* manakins are capable of crossing the river, and there have been a few cases recorded of yellow collared males on the west bank of the river (McDonald et al., 2001; Stein & Uy, 2006b, KML personal observation). Therefore, dispersal ability alone is insufficient to explain why clines under positive selection remain “stuck” at the Río Changuinola, instead of spreading further beyond the river.

Besides dispersal constraints, other mechanisms have been proposed as mediators for the stalling of introgression at the river. For example, Bennett et al. (2021) discusses four hypotheses for the observed phenomenon, including: frequency-dependent selection resulting in differing favored male phenotypes on each side of the river, geographic variation in the color or lighting conditions where males are performing, differences in male choice of display site location for optimal display conspicuousness, and variation in female preference on each side of the river. Presently, however, further research is necessary to understand if any of the proposed mechanisms, or a combination of them, may be playing a role in stalling introgression at the Río Changuinola.

### Melanization as a candidate for selection

Similar to the yellow collar in *M. vitellinus*, belly color introgressed from *M. vitellinus* (dark olive) into *M. candei* (light yellow) and is displaced from the genomic center of the hybrid zone (Brumfield et al., 2001). Here, we find evidence that belly color has continued to introgress and its cline center has moved ∼9 km over the last 30 years, leading to a current phenotypic transition at the Río Changuinola. Given an estimated generation length of ∼2.5 years, averaged from estimates of *M. candei* and *M. vitellinus* from Bird et al. (2020), this means that in about 12 generations belly color has introgressed about 0.7 km per generation. These findings suggest that selection on belly color is ongoing in this hybrid system.

Several non-exclusive factors could explain selection for dark olive bellies in hybrid populations from natural or sexual selection. For natural selection, there could be an ecological or environmental advantage to having darker olive bellies. For sexual selection, both female choice and male-male competition have been implicated as important in lek-breeding systems and the *Manacus* system in particular (McDonald et al., 2001; Stein & Uy, 2006a, 2006b). We propose a few hypotheses for sexual selection: 1) there is a sexual advantage of a yellow collar that is enhanced by contrast with a dark belly, so both introgress together, 2) females prefer dark olive bellies over yellow bellies independently from collar plumage, 3) males with olive bellies are more aggressive and could benefit from male-male competition. We expand on the male-male aggression/melanization hypothesis below.

The olive color in *Manacus* dark bellies is proposed to result from melanin deposition on yellow feathers (Bennett et al., 2021). Pathways controlling aggression and melanin production share some of the same upstream regulators (Ducrest et al. 2008), which has been proposed as an explanation for the observed pattern that animals with darker colorations tend to be more aggressive, especially in birds (Da Silva et al., 2013; de Zwaan, Barnes, & Martin, 2019; Ducrest et al., 2008; Mafli, Wakamatsu, & Roulin, 2011; Mateos-Gonzalez & Senar, 2012). Whether manakins with darker bellies are more aggressive remains to be tested directly, but two previous studies performed aggression assays on *Manacus* manakins based on hybrid status. McDonald et al. (2001) found that hybrid males from the region of yellow plumage introgression were the most aggressive to the presentation of a mount, followed by *M. vitellinus* males, and lastly *M. candei* males. Stein and Uy (2006) performed a similar experiment and found no difference in aggression between yellow and white collared males; however, they worked in a different location along the upper Río Changuinola where the river is narrow and mixed leks with yellow and white collared birds are found.

If aggression in *Manacus* is influenced by melanization (or vice versa), then the difference in response of the manakins tested in the two previous studies could be explained by differences in the introgression of dark melanized bellies. Specifically, McDonald et al. (2001) tested yellow-collared males near sampling site 5, where belly color is typically darker than in leks along the upper Río Changuinola where Stein and Uy (2006) worked. Both studies focused on yellow-collared hybrids, but neither quantified the belly color of the birds tested. If aggression is linked to green bellies/increased melanization more so than yellow collars, their results may be reconcilable. Careful experiments that directly test female preference for olive bellies, male aggression trials with varying olive belly colors, as well as categorizing the genes and metabolic pathways responsible for melanization in this hybrid zone and its causal (or lack thereof) link to aggression, will be necessary to tease apart the connection of introgressing olive bellies, melanization, male-male aggression, and the subsequent effect on hybrid zone dynamics.

Independent of the evolutionary force underlying the movement of belly coloration across the hybrid zone, it is notable that belly color has appeared to have lagged behind collar color. There are several possible explanations for this disparate shift in their movement, including changes in selective pressures (e.g., female choice) over time and/or differences in the genomic architecture of these two coloration traits. First, the traits that females find attractive could change over time or vary within populations (Chaine & Lyon, 2008; DuVal, Fitzpatrick, Hobson, & Servedio, 2023). Within the context of the *Manacus* system, belly color could be under recent selection, while collar color was under stronger sexual selection historically. Another explanation for the lagging introgression of belly color could be due to difference in the genomic architecture between the two traits. For example, the two traits could be the product of independent genomic pathways, as seen in melanin and carotenoid processing genes in *Setophaga* warblers (Baiz, Wood, Brelsford, Lovette, & Toews, 2021). If the loci controlling belly color and collar color are under weak linkage, they would appear to behave as independent units in the genome, and thus move at different rates if under differing selective pressures. Additionally, the genetic architecture underlying both of these traits in *Manacus* is presently unknown, which may influence the movement of a phenotype if the trait is highly polygenic (Sachdeva & Barton, 2018). Interestingly, we observed that belly color has higher variation than collar color in *Manacus*, a pattern also noted by Brumfield et al. (2001). At sites near the Río Changuinola, male belly color exhibits greater within-site variation than collar color, which may indicate that belly color is in fact polygenic in this system. The underlying genomic architecture along with female choice putatively impacting these traits are not yet fully understood and could have important insights into the dynamics of hybrid systems under sexual selection.

### Evolutionary forces and hybrid zone dynamics

The *Manacus* hybrid zone is an ideal system to study the effects of sexual selection, natural selection, and the resulting asymmetrical introgression, in a naturally occurring hybrid zone. In addition to emerging genomic resources, this hybrid zone benefits from multi-year sampling, which is the most reliable way to empirically assess hybrid zone movement (Buggs 2007). Our results indicate that there is an interplay between sexual selection, which leads to the decoupling and movement of certain traits and loci, while other evolutionary forces (such as generalized selection against hybrids) keep the majority of the hybrid genome in relative stasis.

The stability (or lack thereof) of hybrid zones is likely the product of conflicting evolutionary forces. Selection on some traits can be counteracted by generalized selection against hybrids, which in turn produces differential fitness among hybrid individuals and directly affects the movement of alleles and traits across the landscape (Buggs 2007). Sexual selection is one mechanism that can lead to differential reproductive fitness in hybrids. Sexual selection, for example, can reduce hybrid mating success by creating disfavored combinations of traits (Naisbit, Jiggins, & Mallet, 2001; Svedin et al., 2008) or by reducing competitiveness of male sperm (Howard, Gregory, Chu, & Cain, 1998). Yet, despite selection against hybrids being a common outcome of hybridization, sexual selection in hybrid zones can simultaneously fuel the introgression and movement of certain alleles, imparting adaptive potential to new populations (Enard & Petrov, 2018; Pardo-Diaz et al., 2012; Walsh, Kovach, Olsen, Shriver, & Lovette, 2018). Overall, hybrid zones serve as a powerful evolutionary model to study how similar evolutionary forces can have a myriad of outcomes with heterogenous responses across different parts of the genome. Our work here in the *Manacus* hybrid zone demonstrates that traits under sexual selection can move over space and time despite stability in the rest of the genome.

Hybrid zones are ideal cases to observe the dynamics of evolution in action, but evolutionary forces can change over time. Perceived outcomes may shift depending on the population sampled and/or time at which they were sampled, highlighting the importance of following the trajectories of hybrid populations as they evolve (Enbody et al., 2023; Schumer et al., 2017). In this study, we combined sampling over multiple time points and across geographic space, along with both phenotypic and genomic data, to shed light on speciation and selection dynamics in a wild bird. Our results reveal an uncoupling of sexual and natural selection in a natural hybrid zone and evidence for ongoing introgression of dark belly plumage. Despite genomic stability in its center, the *Manacus* hybrid zone is a dynamic system with phenotypes that continue to change over a short timescale. We believe longitudinal sampling in other systems will reveal similar evolutionary dynamics and facilitate deeper understanding of how evolution works in nature.

