## Supplemental Methods and Figures for "Ongoing introgression of a secondary sexual plumage trait in a stable avian hybrid zone"

### Supplemental Information

#### Supplemental Methods

##### Selecting diagnostic parental alleles

To select a panel of diagnostic parental alleles, we first made a subset of the main *Stacks* (Rochette, Rivera-Colón, & Catchen, 2019) catalog to export only genotypes for the parental alleles. This was done by generating a population map file containing only the identifiers for the contemporary *Manacus* individuals from the two putative parental populations, sampling sites 2 and 10. This population map file was then used in the `populations` software (`--popmap`), to first filter for sites present in both parental populations (`--min-populations 2`), present in 80% of individuals per population (`--min-samples-per-pop 0.8`), and with a minimum allele frequency of 5% (`--min-maf 0.05`). In addition, we calculated $F_{ST}$ between the two parental populations (`--fstats`) and deviations from Hardy-Weinberg Equilibrium (HWE) in each population (`--hwe`). This `populations` runs retained 70,311 RAD loci, composed of 367,079 variant sites.

The output of this `populations` run (the `sumstats` and `fstats` output tables) were used as an input of a custom Python program, `make_diagnostic_snp_whitlist.py`. The program first filters the `sumstats` file to retain sites 1) in HWE in both parental populations (`--hwe`, `--hwe-alpha=0.05`) and 2) with a minimum minor allele frequency greater than 5% in either population (`--min-maf=0.05`). The `fstats` file is then processed to retain only sites with an $F_{ST}$ above 0.8 between both populations (`--min-fst=0.8`), maximizing the differences in allele frequency in each population at each observed site. The overlap of both sets contains the pool of sites available post-filtering (3,844 sites total), which we then filtered to export only one site per RAD locus to retain independence between makers (`--write-random-snp`) and exporting a total of 2,000 sites (`--n-sites=2000`). The output of the program is a *Stacks* catalog whitelist containing the ID of the retained sites, which represent the subset of diagnostic markers between parental populations. This new whitelist was then used as an input for a second run of the `populations` software (`--whitelist`), along with the parental population map file (`--popmap`), in order to export the genotypes for the target parental diagnostic sites into a VCF file (`--vcf`, `--ordered-export`).

##### Simulation of hybrid genotypes

The custom Python program `sim_hybrids_from_parents_vcf.py` was used to simulate genotypes for multi-generation hybrid individuals generated from two parental populations ($P_1$ and $P_2$). The script takes a set of previously processed set of diagnostic parental genotypes across multiple genomic sites and begins by calculating the observed allele frequency at each site for each genomic location. These frequencies are stored in a parental allele dictionary, which represents the pool of parental alleles from which new hybrid individual genotypes will be sampled in subsequent generations (i.e., at site $x$, the allele frequencies are $f(p) = 0.8$ for $P_1$ and $f(p) = 0.2$ for $P_2$). At this stage, the script also retains the genomic information for each site in the VCF (e.g., chromosome and base pair coordinates) for downstream compatibility.

Next, the script defines a set of hybrid crosses, specifying the different types of hybrid to simulate. Each cross is represented by a class defining the generations ($g$) of hybridization observed in each hybrid type (e.g., $g = 1$ for $F_1$ hybrids, $g = 2$ for $F_2$, etc.), as well as the corresponding parents of each cross (e.g., the two parents of an $F_1$ cross are one each from $P_1$ and $P_2$, the parents of an $F_2$ are both from the $F_1$ cross, etc.). The cross class also specifies the information of the backcross individuals, such as their generation and the direction of the backcross to either $P_1$ or $P_2$. Except for generations $g = 0$ (composed of pure parental $P_1$ or $P_2$ individuals) and $g = 1$ (composed of $F_1$ individuals), three crosses are generated per generation: 1) a hybrid x hybrid cross originating from the cross of two hybrid individuals from the previous generation ($g - 1$), 2) a hybrid x $P_1$ backcross, originating from $g - 1$ generations of crossing against $P_1$, and 3) a hybrid x $P_2$, with $g - 1$ crossing generations against $P_2$. We limit to these three crosses per-generation in order to limit the possible number of parental combinations per cross in later generations. Additionally, the cross class specifies the parental genomic proportions expected in each cross. As specified by (Fitzpatrick, 2012; Turelli & Orr, 2000) these three proportions are 1) the proportion of sites with both alleles originating from $P_1$ ($p_{11}$), 2) the proportion of sites with one allele each from $P_1$ and $P_2$ ($p_{12}$; also defined as the interspecific heterozygosity), and 3) the proportion of sites with both alleles from $P_2$ ($p_{22}$). For example, the genomic proportions of an $F_1$ hybrid individual are specified as $p_{11} = 0$, $p_{12} = 1$, and $p_{12} = 0$, as all their loci contain one copy each of both parental alleles, indicating a one-to-one admixture of both parental genomes. Each hybrid cross (and backcross) thus contains different genomic proportions based on the expected degree of parental admixture. These proportions are known for early-generation hybrids (e.g., see Table 1 in (Fitzpatrick, 2012)), but can additionally be determined based on the generations of hybridization and the type of cross (Fitzpatrick, 2008; Lynch, 1991) and thus can be calculated for late-generation hybrids and backcrosses.

After crosses are determined, the expected genotype frequencies for each specified cross are determined for each sampled locus. For each cross type, the expected genotype frequencies are calculated based on the parental allele frequencies at the given locus and the expected genomic proportions of each cross. Using formula 1 from (Fitzpatrick, 2012), the probability of being homozygous for allele $p$ at a given locus can be defined as:

$$f(p,p) = p_{11}{p_{P_1}}^2 + p_{12}p_{P_1}p_{P_2} + p_{22}{p_{P_2}}^2$$

where, $p_{11}$, $p_{12}$, and $p_{22}$ represent the three genomic proportions, while $p_{P_1}$and $p_{P_2}$represent the observed frequencies of allele $p$ in $P_1$ and $P_2$, respectively. Similarly, the probability of being heterozygous for alleles $p$ and $q$ can be defined as (formula 2 from (Fitzpatrick, 2012)):

$$f(p,q) = p_{11}{}^2 p_{P_1} q_{P_1} + p_{12}\left(p_{P_1} q_{P_2} + q_{P_1} p_{P_2}\right) + p_{22}{}^2 p_{P_2} q_{P_2}$$

where $q_{P_1}$and $q_{P_2}$represent the observed frequencies of allele $q$ in $P_1$ and $P_2$, respectively. We use these probabilities to sample an individual genotype at the specified locus. Since we assume independence between the provided parental makers, genotypes are independently sampled at each locus. The final genotype matrix, containing the simulated genotypes for each sample at each locus is then stored in VCF format.

We ran the `sim_hybrids_from_parents_vcf.py` simulation script on the set of 2,000 diagnostic makers (described above) between *M. vitellinus* and *M. candei*. We specified the assignment of parental individuals as $P_1$ and $P_2$ using a population map file (`--popmap`), specifying the previously generated diagnostic loci VCF as input (`--vcf`). To generate both early- and late-stage hybrids, we simulated 10 generations of hybridization and backcross (`--generations=10`), simulating 50 individuals per cross (`--n-individuals=50`). In total, we simulated genotypes for 1,500 individuals across 30 different crosses (including hybrid x hybrid crosses, backcrosses to both $P_1$ and $P_2$, as well as pure $P_1$ and $P_2$ individuals).

#### Filtering schemes for genetic geographic cline centers

As described in the methods, out base filtering scheme for the genetic geographic clines is based on the merging of the Historical and Contemporary datasets. For each of the two datasets, we first remove SNPs that are either missing or that were identified as non-clinal based on the assignment of a "null" cline model by *HZAR* (Derryberry, Derryberry, Maley, & Brumfield, 2014). Secondly, we removed SNPs for which the cline center estimates lie beyond the sampled boundaries of the hybrid zone (outside the range of 0 and 92.5km, based on the locations of sampling site 10 and 2, respectively). We kept only SNPs that were present in both Contemporary and Historical datasets after applying these filters. This is our baseline-level filtering (referred hereafter as "base filters").

Next, in addition to the base filters described above, we applied a second filter based on the difference between the observed parental allele frequency between the two parental sampling sites (sites 2 and 10). The aim of this filter is to retain only a subset of SNPs which maximize the observe difference (i.e., are diagnostic) between the two parental populations. For each dataset, we removed SNPs for which the difference between parental allele frequencies was below a specified threshold. We only retained SNPs that were present in both Contemporary and Historical datasets after applying the base and parental allele frequency filters. We applied these filters at three levels of stringency, removing SNPs for which the parental allele frequencies differed by less than 80%, 50%, 20%, and 10%. These filtering schemes are notated as “parental 80%”, “parental 50%”, “parental 20%”, and “parental 10%”, respectively.

Lastly, we applied a third filtering scheme based on the range of the Bayesian credibility interval (CI) of the estimated best fit clines estimated by *HZAR* (referred to as “CI filter”). The aim of this filtering scheme was to only retain clines with narrower CI, as a proxy for the accuracy of the cline center estimation. In addition to the base filter, for each dataset, we removed SNPs when their estimated cline center CI interval was above a specified threshold, keeping only SNPs where clines for both historical and contemporary datasets were retained after applying these filters. We selected a CI filtering threshold of 11.56 Km (the mean inter-sampling site distance), with the intention of removing cline centers for which the CI range was larger than the distance between sampling locations. Summary statistics for all 5 filtering schemes are reported in Table S7.

#### Classifying orthologous chromosomes in manakins

We used a conserved synteny analysis to assign orthology between the reference assemblies of the golden-collared manakin *M. vitellinus* (NCBI RefSeq: GCF_001715985.3) and the chromosome-level assembly of the lance-tailed manakin *Chiroxiphia lanceolata* (NCBI RefSeq: GCF_009829145.1). This analysis of conserved synteny was performed using the *Synolog* software (Catchen, Conery, & Postlethwait, 2009; Small et al., 2016). First, we performed reciprocal *BLAST*s of the annotated protein sequences of both genomes using the `blastp` algorithm from *BLAST*+ v2.4.0. Next, we input these *BLAST* hits into *Synolog* along with the genome annotation coordinates (GTF) for each assembly. *Synolog* then identifies orthologous genes through reciprocal best *BLAST* hits and uses the genome coordinates to find their location in their respective genome. Once the location of the orthologs is determined, *Synolog* builds clusters of conserved synteny between local gene neighborhoods, which are expanded to determine orthology across chromosomes/scaffold sequences. Based on these patterns of conserved synteny, we assigned each scaffold in the *M. vitellinus* assembly to its corresponding chromosome-level ortholog in *C. lanceolata.*

#### Supplemental Figures

##### Figure S1.

Scatterplot of the mean percent reflectance of belly color for every male manakin used in the study. The USB2000 spectrophotometer (in green) was used in the 2017 field season, while the Flame spectrophotometer (in purple) was used both in the field in 2018-2020, and on the museum skins in 2022. The location is denoted by the sampling site ID from site 10 to 2. While all historical specimens had reads taken on the same spectrophotometer, some of the contemporary, wild-caught spectrophotometry reads were taken entirely on the Flame or entirely on the USB2000 spectrophotometers (e.g., sampling site 6 had all spectrophotometry reads taken on the USB2000 spectrophotometer). Note that all reads taken in sampling sites 3 and 4 (on each side of the Río Changuinola) were taken on the same spectrophotometer, which are the same locations we detected the largest changes over time in belly color. Overall, reads taken with these two different spectrophotometers roughly overlap each other. Differences in spectrophotometry reads appear to correlate with the temporal dataset rather than the spectrophotometer used. See Table S1 for summary statistics of the mean, median, and standard deviation of for each spectrophotometer, temporal dataset, and sampling site.

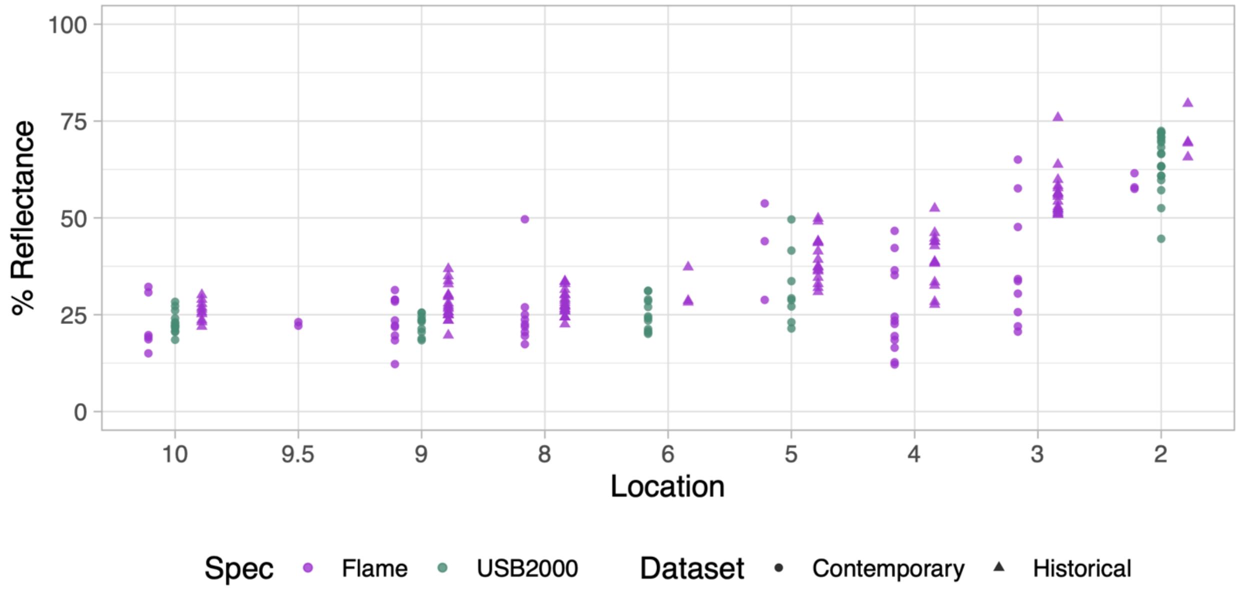

**Figure S2.**

Histogram of all cline centers for all 48,316 clinal SNPs without (A) and with (B) population 9.5 in the contemporary dataset to visualize the raw SNP cline data. The dashed lines represent the median cline center for the contemporary and historical populations in their respective colors. The blue dotted line is the location of the Río Changuinola. Note that all 48K clines are plotted here and some clines have credibility intervals that are quite large (as large as the whole hybrid zone sampling space). Overall see that the histograms have similar patterns through time and nearly identical median cline centers.

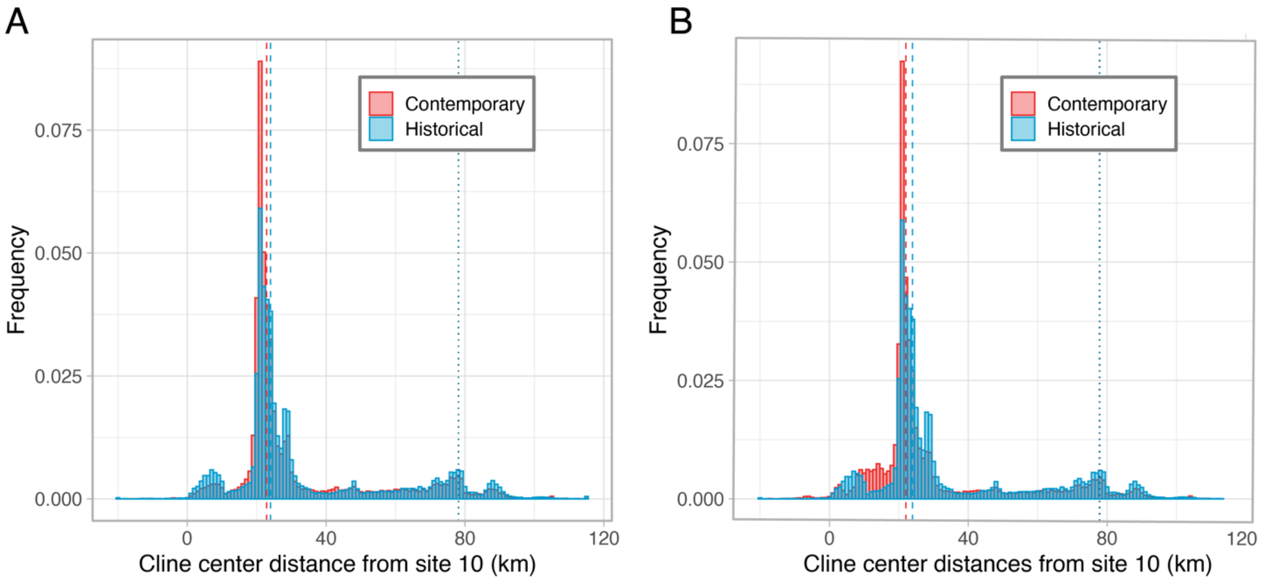

**Figure S3.**

Boxplots showing the distribution of genomic hybrid index values of historical and contemporary transect samples. A hybrid index of 0 indicates complete *M. candei* ancestry (site 2) while a blue of 1 indicates complete *M. vitellinus* ancestry (site 10). Blue boxplots are the historical hybrid indexes and red are the contemporary. The diamond shape is the mean hybrid index for that sampling site. Note that there is no blue box for sampling site 9.5, as it was first characterized in this study and has no historical counterpart.

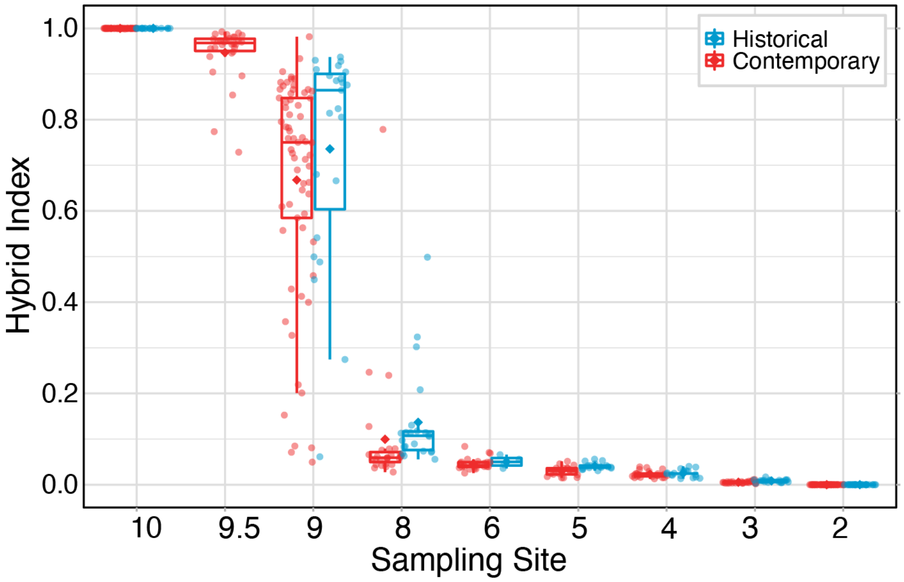

**Figure S4.**

Hybrid classification “Triangle” plots showing the relationship between genomic hybrid index and interspecific heterozygosity (proportion of sites with alleles from both parental populations) for parental and hybrid individuals. A) Hybrid classification of the empirical *Manacus* data using 2,000 diagnostic SNPs. Colors denote the sampling locations along the hybrid zone, while the shape denotes the two temporal datasets. We do not find evidence of any early generation (F1 or F2) hybrids. B) Classification of simulated hybrid crosses based on 2,000 diagnostic SNPs derived from our empirical parental *M. candei* and *M. vitellinus* populations. Colors denote the generation of hybridization (1 generation for F1, 2 generations for F2, etc.), while the shape denotes the type of cross (i.e., crosses between two parents, crosses between two hybrids, or backcrosses with either parent). The hybrid classification from the simulated data serves as a comparison to the empirical data to further support the absence of early generation hybrids or recent backcrosses in the *Manacus* hybrid zone.

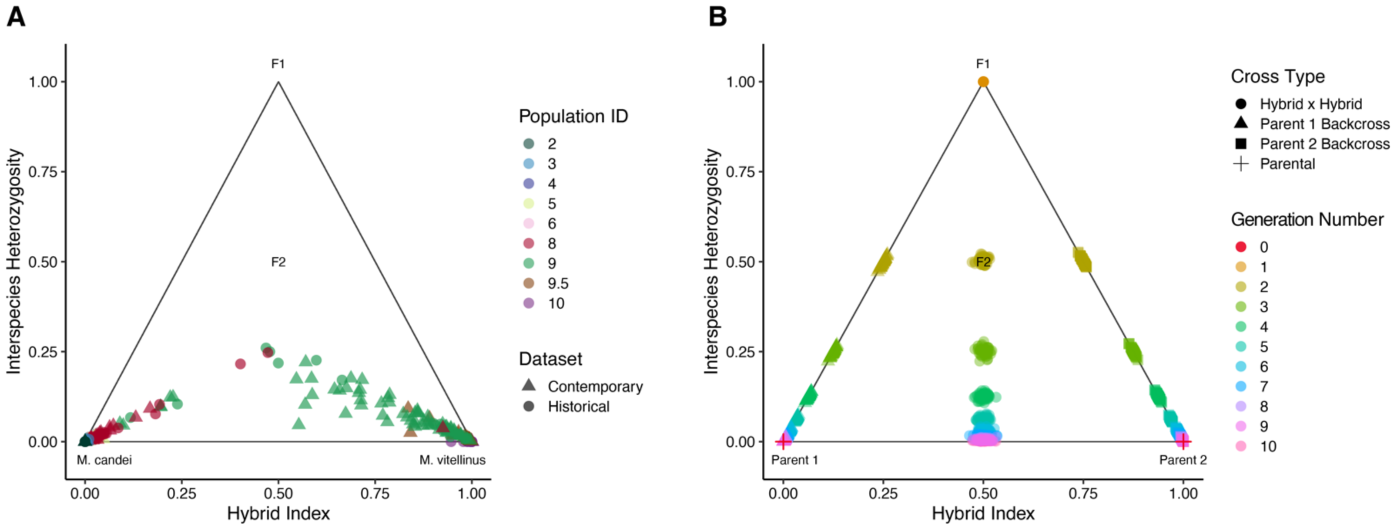

#### Supplemental Tables

[See attached Excel document]

##### Table S1.

Summary statistics for belly color measurements taken on each spectrophotometer in both temporal datasets.

##### Table S2.

Sample metadata for the genetic samples sequenced. Sample index is a sortable index for the table. Sample ID for contemporary birds was based on the bird's aluminum band number. The Sample ID for the Historical birds was based on deprecated USNM tissue ID numbers.

##### Table S3.

Library specific metadata for *Manacus* RADseq genomic sequencing data.

##### Table S4.

CV Errors from the *ADMIXTURE* analysis and corresponding genetic clustering (K) values.

##### Table S5.

Phenotypic and hybrid index cline center and width estimates with associated 95% credibility intervals (CI) for the historical and contemporary datasets.

##### Table S6.

Mean, median, and standard deviation of the genomic hybrid index of the historical and contemporary datasets by sampling site. Sample sizes match those presented in Table 1.

##### Table S7.

Summary statistics for the genetic (SNP) geographic clines within and between historical and contemporary datasets.

##### Table S8.

Distribution of clines with detectable movement across the *Manacus vitellinus* reference assembly (GCF_001715985.3) of the base and 80% parental diagnostic SNP filtering schemes. Chromosome number assignments are from orthologous sequences in the *Chiroxiphia lancaeolata* chromosome-level reference assembly (GCF_009829145.1).

**Table S9.**

Recorded values for the male phenotypic plumage data used in the geographic cline analyses. The Genetic Sample ID # matches the Sample ID # in Table S1.

**Table S10.**

Summary of plumage measurements across the hybrid zone. Mean, standard deviation (SD), and number of individuals measured (n) are presented for each sampling site and historic vs contemporary dataset. Sample site 9.5 was added in the contemporary dataset.
